# Sustained attention and spatial attention distinctly influence long-term memory encoding

**DOI:** 10.1101/2020.09.14.297341

**Authors:** Megan T. deBettencourt, Stephanie D. Williams, Edward K. Vogel, Edward Awh

**Affiliations:** Institute for Mind and Biology, University of Chicago, Chicago, IL 60637; Department of Psychology, University of Chicago, Chicago, IL 60637; Department of Psychological & Brain Sciences, Boston University, Boston, MA, 02215; Grossman Institute for Neuroscience, Quantitative Biology and Human Behavior, University of Chicago, Chicago, IL 60637

**Keywords:** top-down attention, working memory, multivariate pattern analysis

## Abstract

Our attention is critically important for what we remember. Prior measures of the relationship between attention and memory, however, have largely treated “attention” as a monolith. Here, across three experiments, we provide evidence for two dissociable aspects of attention that influence encoding into long-term memory. Using spatial cues together with a sensitive continuous report procedure, we find that long-term memory response error is affected by both trial-by-trial fluctuations of sustained attention and prioritization via covert spatial attention. Furthermore, using multivariate analyses of EEG, we track both sustained attention and spatial attention prior to stimulus onset. Intriguingly, even during moments of low sustained attention, there is no decline in the representation of the spatially attended location, showing that these two aspects of attention have robust but independent effects on long term memory encoding. Finally, sustained and spatial attention predicted distinct variance in long-term memory performance across individuals. That is, the relationship between attention and long-term memory suggests a composite model, wherein distinct attentional subcomponents influence encoding into long-term memory. These results point towards a taxonomy of the distinct attentional processes that constrain our memories.

## Introduction

In our daily lives, we fail to remember many of the items that we encounter, largely because we were not paying sufficient attention. While attention and long-term memory are clearly intertwined (Aly & Turk-Browne, 2017; Chun & Turk-Browne, 2007), past work investigating the relationship between attention and long-term memory often considers attention to be a monolithic cognitive construct. However, attention has numerous subcomponents, any one of which could underlie the relationship with memory (Chun et al., 2011; J. Fan et al., 2005; Hakim et al., 2019; Poole & Kane, 2009; Robison & Brewer, 2019). For example, sustained attention fluctuates from trial to trial, and spatial attention can be oriented to different locations in space. However, studies investigating long-term memory have generally lacked the ability to disentangle whether memory failures reflect poor sustained attention, misallocated spatial attention, or both. This raises two potential explanations for the relationship between attention and long-term memory: In a *unified* model of attention and memory, memory failures could be attributable to convergent failures of multiple forms of attention. That is, lapses of sustained attention coincide with moments when spatial attention is oriented to the wrong location and together influence memory encoding. Alternatively, in a *composite* model of attention and memory, memory failures arise from failures of any particular attentional subcomponent. That is, lapses of sustained attention and failures of spatial attention could each independently account for memory encoding. Thus, the goal of this study was to determine whether sustained attention and spatial attention exert overlapping or distinct influences on long-term memory.

Sustained attention and spatial attention are each critically important for long-term memory. Sustained attention fluctuates between advantageous and disadvantageous states over time. These trial-by-trial fluctuations of sustained attention predict which items are later remembered (deBettencourt et al., 2018). In fact, fluctuations of sustained attention remain a key candidate explanation for previous demonstrations of pre-trial states that predict long-term memory (Gruber & Otten, 2010; Guderian et al., 2009; Otten et al., 2006; Weidemann & Kahana, 2020). Sustained attentional state can be monitored via behavioral performance and via multivariate analyses of brain data (deBettencourt et al., 2015; Esterman et al., 2013; Rosenberg et al., 2016). On the other hand, spatially cueing an item also improves long-term memory (LaRocque et al., 2015; Turk-Browne et al., 2013; Uncapher et al., 2011; Ziman et al., 2019). The influence of spatial cues can be measured behaviorally and via multivariate analyses of brain data (Foster et al., 2017; Sprague & Serences, 2013). A critical challenge is to understand how sustained attention and spatial attention interact and give rise to long-term memories.

In this study, we examine whether sustained and spatial attention exert distinct or common influences on long-term memory encoding using behavioral and neural signatures. To measure long-term memory behavior with high sensitivity, we employed a continuous report task, in which participants report a particular dimension of a stimulus along a continuous space (Biderman et al., 2019; J. E. Fan & Turk-Browne, 2016; Richter et al., 2016; Sutterer & Awh, 2016; Tompary et al., 2020; Xie et al., 2020). To resolve the moment-by-moment influence of attention on long-term memory, we analyze multivariate, multi-frequency EEG signals during time intervals prior to encoding. In Experiment 1, we present a behavioral paradigm that captures how sustained and spatial attention distinctly influence long-term location memory on a continuous report task. In Experiment 2, we identify a multivariate EEG signature of sustained attention that predicts performance independent of variations in spatial attention. In Experiment 3, we extend these findings to show that spatial and sustained attention influence color memory. Finally, collapsing across all studies, we show that individual differences in sustained and spatial attention predict unique variance in long-term memory performance.

## Experiment 1

The goal of this experiment was to examine whether signatures of sustained and spatial attention predict long-term memory. We hypothesized that long-term memory would reflect trial-by-trial fluctuations of sustained attention, as well as the prioritization of cued stimuli by spatial attention. We obtained a sensitive measure of long-term memory accuracy by asking participants to report their memory for the spatial location of trial-unique objects using a continuous report task.

### Methods

#### Participants

In Experiments 1a and 1b, a combined total of fifty-two adults participated for University of Chicago course credit or $20 payment ($10/hour). In all studies we targeted data collection from twenty-five participants prior to exclusion. In Experiment 1a, twenty-five adults (15 female, mean age=23.2 years) participated, and in Experiment 1b twenty-seven adults participated (17 female, mean age=24.2 years). We excluded any participants whose performance exceeded three standard deviations from the population mean (*n*=2 in Experiment 1a; *n*=1 in Experiment 1b) and participants who were outliers in terms of study completion (*n*=2 in Experiment 1b completed 50% of the study in the allotted time). Therefore, the final sample of participants was 23 for Experiment 1a and 24 for Experiment 1b. All participants in this experiment and the following experiments reported normal or corrected-to-normal color vision and provided informed consent to a protocol approved by the University of Chicago IRB.

#### Apparatus

Participants were seated facing a LCD monitor (120-Hz refresh rate) in a testing room. In Experiment 1a, participants were approximately 70 cm from the monitor and in Experiment 1b they were approximately 88 cm from the monitor, due to a reconfiguration of the behavioral testing rooms. Stimuli were presented in Python using Psychopy (Peirce, 2007).

#### Stimuli

Trial-unique real-world object pictures were presented on a gray background (Brady et al., 2008; Brodeur et al., 2014). At encoding, these images (subtending 3° visual angle) were presented along a light gray ring (at 5° eccentricity). A black fixation dot (0.5°) appeared at the center of the screen. Peripheral spatial cues (black dots, 0.5°) appeared along the gray ring.

#### Procedure

In Experiment 1a, on each working memory trial, a peripheral spatial cue briefly appeared (250 ms) along the ring and participants were instructed to covertly attend to the cued spatial location (Figure **1a**). After an extended pre-stimulus delay (2–4 s), four items (trial-unique object pictures) briefly appeared along the ring (250 ms). Participants were instructed to hold the items in mind over a retention interval (2 s). Then, one of the items reappeared at the center, and the mouse cursor was initialized to a random position along the ring. Participants reported the original location of the probed item by clicking along the ring with the mouse. On validly cued trials (75%), the item that re-appeared had been presented at the cued location (*cued*). On invalidly cued trials (25%), one of the other 3 uncued items was selected (*uncued*). After the response, there was a blank inter-trial interval (1–2 s). Participants were instructed to maintain central fixation throughout the trial. Participants completed 16 blocks, each consisting of 24 working memory trials, and the cue position was counterbalanced across trials within a block.

**Figure 1.**
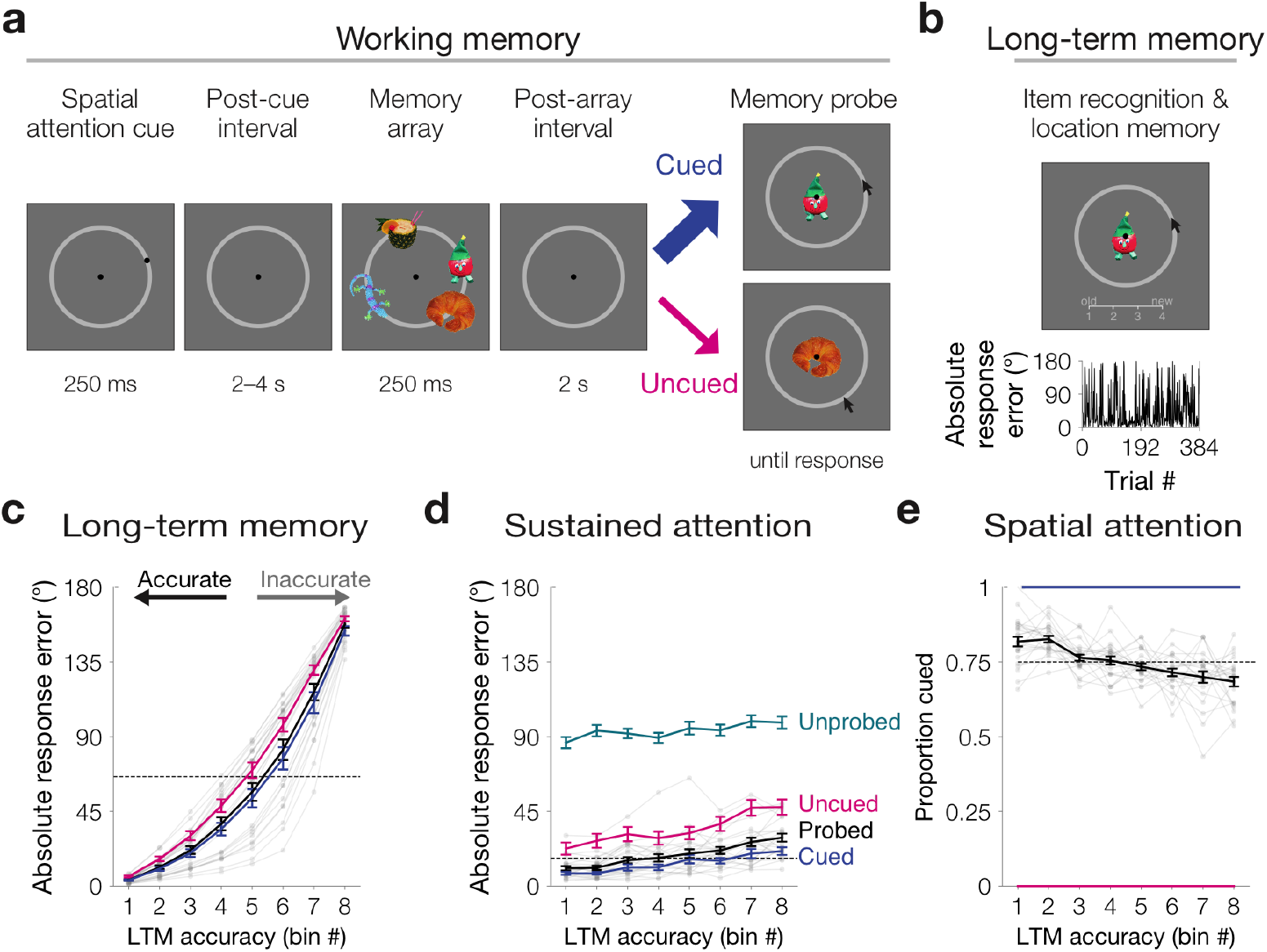
Experiment 1a results. **a** In the working memory phase, each trial was comprised of a brief peripheral spatial cue, a blank pre-stimulus delay, a memory array of four real-world trialunique object pictures, and a blank retention interval. One of the items (75% cued item, 25% uncued item) reappeared at the center, and participants reported its original location by clicking along the ring. **b** In the long-term memory phase, participants reported their long-term recognition memory and location memory (upper). Long-term memory accuracy was quantified as absolute response error (0–180°), which fluctuated substantially across trials (lower). **c** Long-term memory variability binned across trials. Location memory absolute response error (0–180°) was sorted and binned into octiles within participant for all probed items (black line). To equate for whether an item was cued to be attended, we also repeated all analyses separately for cued items (blue), and uncued items (pink). The median long-term memory response error for all probed items is depicted as a dashed black line. **d** Memory outcomes reflect trial-by-trial fluctuations of sustained attention. Memory encoding success was operationalized as absolute response error (0–180°) for items within each bin. For uncued (pink), probed (black), and cued (blue) items, absolute response error was obtained during the working memory phase The median working memory response error for probed items is depicted as a dashed black line. Unprobed items (teal) were, by definition, not tested in the working memory phase. Therefore, absolute response error for these items was obtained during the long-term memory phase (0–180°). The slope of each line is positive across the bins (*p*s<0.05). **e** Memory outcomes reflect trial-by-trial differences of spatial attention. In each bin, we calculated the proportion of trials that had been cued. For all probed items (black), The proportion of trials that were cued decreases across the eight bins (*p*<0.001). The dashed line represents the mean proportion cued (0.75). Cued items (blue) and uncued items (pink) controlled for whether an item was cued to be attended (100% and 0% cued, respectively). Error bars depict the standard error of the mean. Data from each participant for all probed items are overlaid in small gray dots connected with lines.

In the long-term memory phase, all probed items reappeared (24 per block) along with all items that had cued but not probed from the uncued trials (6 per block). A balanced number of new object pictures appeared (30 per block); these items otherwise never appeared during the experiment. First, participants completed a recognition memory rating for each item. Below the image, a four-point confidence rating scale appeared below the image. When participants made their responses (using the keys 1–4), it was briefly displayed on the scale (0.5 s). Then, for all old items, participants completed a location memory continuous report. The fixation dot turned white and the participant was instructed to retrieve the original location of that item for 1 s. Then, the fixation dot turned black, the mouse cursor appeared randomly along the ring, and the participant reported the original location for that object along the ring. After each item, there was a blank inter-trial interval (0.5 s).

In Experiment 1b, half of the blocks were identical to Experiment 1a with a single cue that was either valid or invalid (*cued/uncued*). In the other half of the blocks, the encoding arrays were preceded by four dots (*neutral*). These four dots indicated the location of each of the items, but provided no information about which item was most likely to be tested. As such, this condition controlled for the visual presentation of peripheral dots and provided the same temporal information about an upcoming memory array (Jonides & Mack, 1984). The order of single cued blocks and neutral cued blocks were randomized, such that there were two of each type every four blocks.

Participants performed this task for 2 hours or until they completed 16 blocks in total (384 trials). In Experiment 1a, participants completed on average 345 (89.95 %) of the maximum 384 trials, ranging from 240–384. In Experiment 1b, participants completed 360 (93.75 %) of the maximum 384 trials, ranging from 312–384.

#### Behavioral analysis

Location memory was measured via continuous report and analyzed as response error, or the angular difference between the original minus reported location (−180° to 180°). For recognition memory, high-confidence old responses were treated as remembered and all other responses as forgotten to calculate item recognition memory hit rate. Recognition memory performance was summarized as a single nonparametric measure of sensitivity (*A′*).

#### Statistics

Descriptive statistics are reported as the mean and 95% confidence interval (CI) of the bootstrapped distribution. If the hypothesis was directional, one-sided tests were performed. Because some of the data violated the assumption of normality, nonparametric statistics were performed by resampling participants with replacement 100,000 times. The *p*-value corresponds to the proportion of the iterations in which the bootstrapped mean was in the opposite direction (Efron & Tibshirani, 1986). Any *p*-values smaller than one in one thousand were approximated as *p*<0.001.

### Results

#### Experiment 1a

We obtained a sensitive measure of long-term location memory as the absolute response error from the original location (*d*_ltm_=61.00°, 95% CIs 54.36–66.83; Figure **1b**). We conducted a subsequent memory analysis, sorting trials from each participant into eight bins (Figure **1c**). The mean absolute response error per bin spanned a wide range of performance, from the most accurate bin (*d*_1_=3.65°, 2.97−4.43°) to the least accurate bin (*d*_8_=158°, 153–161°).

We hypothesized that long-term memory would relate to performance during the working memory task. We examined response error in the working memory phase as a measure of encoding quality (*d*_wm_=18.86°, 95% CIs 16.11–23.17). We calculated a linear fit to relate longterm memory performance (i.e., bin number) to working memory performance (i.e., response error). If long-term memory related to working memory, then working memory response error would be larger for the worst remembered items. Indeed, we observed a reliably positive slope relating long-term memory to working memory (*m*=2.71, 2.11–3.46; one-tailed p<0.001; Figure **1d**). This analysis revealed a general correlation between memory encoding and long-term memory. This is consistent with prior work that has found that better working memory serves as a more efficient “gateway” for long-term memory (Fukuda & Vogel, 2019).

The correlation between working memory and long-term memory could be explained in two distinct ways. On the one hand, successful prioritization of the cued item over the other items in the display (*spatial attention*) could influence both working memory and long-term memory. On the other hand, memory outcomes might reflect a broader fluctuation of attentional state (*sustained attention*) that could impact the quality of memory for all items, regardless of cueing. We examined each of these factors in turn:

First, we examined the influence of spatial attention on long-term memory. We predicted that spatial cues would benefit long-term memory. Indeed, items that were cued to be spatially attended were better remembered in the long-term memory phase (*d*_cued_=57.95°, 50.69–64.40°; *d*_uncued_=70.14°, 65.03–74.78°; one-tailed *p*<0.001). We also examined the influence of spatial attention on the long-term memory bins. We calculated a linear fit to relate long-term memory bins to the proportion of trials that had been cued in each bin. We predicted that there would be more cued items in the most accurate bins. Indeed, we observed a reliably negative slope across bins (*m*=−0.02, −.03–−0.01; one-tailed *p*<0.001; Figure **1e**). Thus, spatially attending to items enhanced long-term memory fidelity.

Next, we examined the influence of sustained attention on long-term memory. We predicted that long-term memory encoding would be affected by trial-by-trial fluctuations of sustained attention. To control for spatial attention, we repeated the binning analysis separately for *cued* and *uncued* items (Figure **1c&d**). We observed a positive slope between long-term memory and working memory response error for cued items (*m*=2.02, 1.49–2.78; one-tailed *p*<0.001) and uncued items (*m*=3.53, 2.24–4.72; one-tailed *p*<0.001). This revealed that trial-by-trial fluctuations appeared to influence long-term memory encoding, regardless of whether the items were spatially attended.

We further posited that fluctuations of sustained attention could broadly impact memory for multiple items from the same display. That is, sustained attention could operate in an item-general fashion, reflecting a common mechanism by which memory for all simultaneously displayed items fluctuated synchronously. Alternatively, sustained attention could reflect item-specific fluctuations; for example, reflecting memory success only at covert spatial orienting. We generally measured long-term memory for a single item per display, which is unable to differentiate between these two accounts. However, on trials when the cue was invalid, we also measured long-term location memory of the item that was initially cued but unprobed. Overall, long-term memory for these unprobed items (*d*=93.31°, 90.79–95.61°) was worse than memory for the items probed in the working memory phase, reflecting a testing effect (one-tailed *p*<0.001). Since we did not obtain a working memory response for these items, we examined whether the long-term memory for the uncued item related to the long-term memory for the unprobed item. Indeed, there was a positive slope relating the long-term memory bin (i.e., location memory of the uncued item) to long-term memory for the unprobed item (*m*=1.49, 0.64– 2.49; one-tailed *p*<0.001; Figure **1d**). This provides evidence that sustained attention fluctuations broadly impact memory for multiple items from the same display. This further implicates trial-by-trial fluctuations of sustained attention, rather than fluctuations of the degree to which observers prioritized the cued location.

#### Experiment 1b

We conducted this experiment to replicate and extend the previous findings, by comparing performance with single cues (valid or invalid; Figure **2a**). Importantly, in half of the blocks, all images were preceded by neutral cues (Figure **2b**). In the neutral condition, the four cues indicated the spatial location of each item but provided no information about which item was likely to be tested. This enabled us to control for any location information a spatial cue may have provided. We again operationalized location memory performance as absolute response error (*d*=63.73°, 58.43–68.52°) and repeated the binning analysis (Figure **2c**). We hypothesized that long-term memory was linked with attention at encoding, which would replicate our findings from Experiment 1a. We measured encoding success using the behavioral measure of response error from the working memory phase (*d*=23.23°, 19.79–26.90°). Indeed, we again observed a reliably positive slope relating long-term memory (i.e., bin number) to working memory response error across all probed items (*m*=3.54, 2.84–4.44; one-tailed p<0.001; Figure **2d**). This replicated the finding of a general correlation between memory encoding and long-term memory.

**Figure 2.**
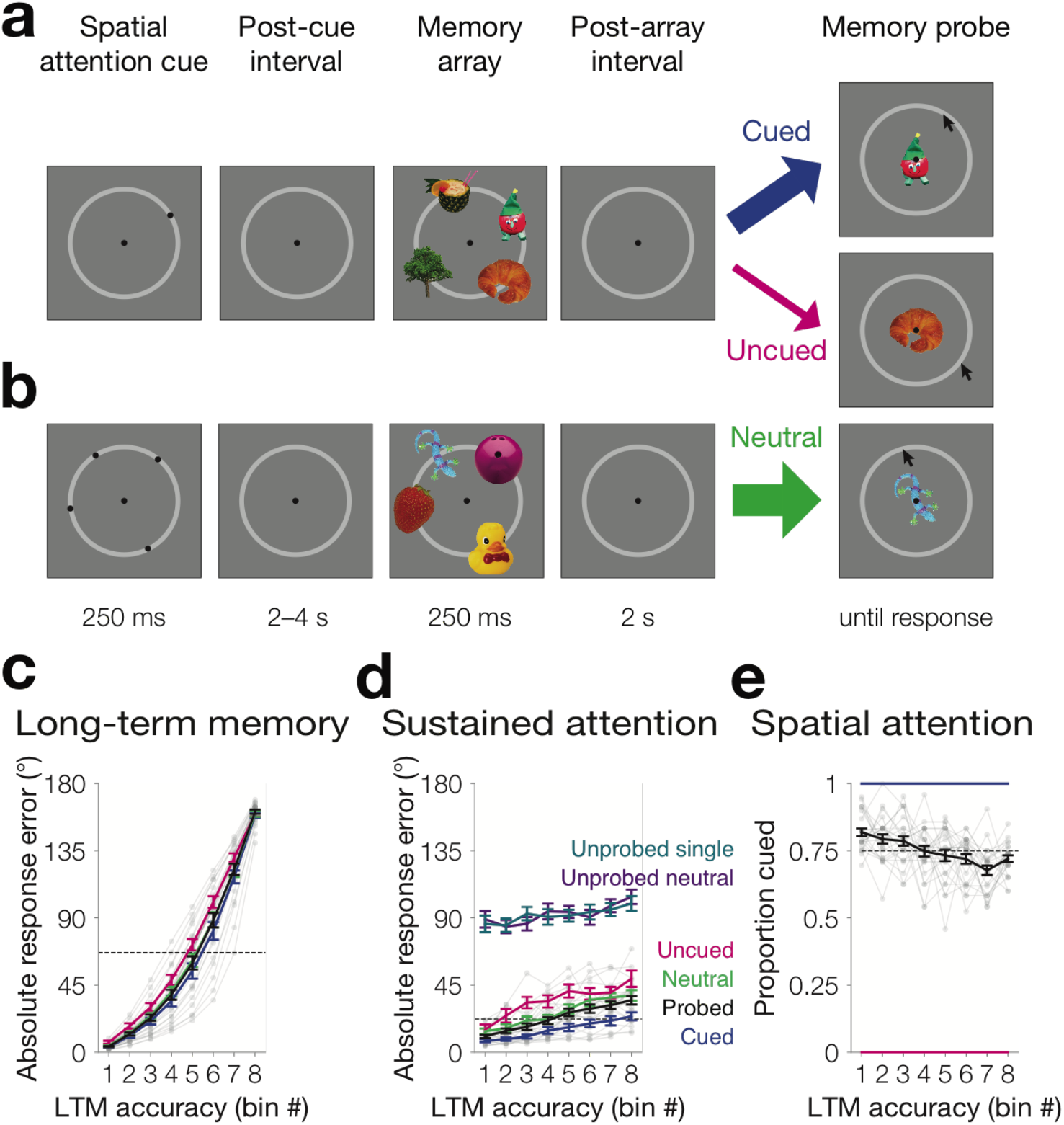
Experiment 1b results. **a** Task design for the single-cued condition. Half of the blocks were the same as Experiment 1a. **b** Task design for the neutral condition. In half of the blocks, all four items were preceded by a spatial cue. **c** Long-term location memory variability binned across trials. Absolute response error (0–180°) was sorted and binned into octiles within participant for all probed items (black line). To equate for whether an item was cued to be attended, we also repeated all analyses separately for cued items (blue), uncued items (pink), and neutral items (green). The median long-term memory response error for all probed items is depicted as a dashed black line. **d** Memory outcomes reflect trial-by-trial fluctuations of sustained attention. Memory encoding success was operationalized as absolute response error (0–180°) for items within each bin. For uncued (pink), neutral (green), probed (black), and cued (blue) items, absolute response error was obtained during the working memory phase The median working memory response error for probed items is depicted as a dashed black line. Unprobed items from the single cued bocks (teal) and the neutral cued blocks (purple) were not tested in the working memory phase. Therefore, absolute response error for unprobed items was obtained during the long-term memory phase (0–180°). The slope of each line is positive across the bins (*p*s<0.05). **e** Memory outcomes reflect trial-by-trial differences of spatial attention. In each bin, we calculated the proportion of trials that had been cued. For all probed items (black), the proportion of trials that were cued decreases across the eight bins (*p*<0.001). The dashed line represents the mean proportion cued (0.75). Cued items (blue) and uncued items (pink) controlled for whether an item was cued to be attended (100% and 0% cued, respectively). Error bars depict the standard error of the mean. Data from each participant for all probed items are overlaid in small gray dots connected with lines.

We next examined whether this general correlation reflected distinct contributions of spatial and sustained attention. Indeed, spatial attention influenced long-term memory: Neutrally cued items were remembered worse than cued items (*d*_cued_=60.39°, 54.60–65.78; *d*_neutral_=64.32°, 58.66–69.27; one-tailed *p*<0.001) and better than uncued items (*d*_uncued_=71.12°, 66.87–75.82; one-tailed *p*=0.004). Moreover, we examined the effect of spatial attention within the long-term memory bins, by replicating our analyses from Experiment 1a within the single cued blocks. We replicated the finding from Experiment 1a that the proportion cued decreased across bins (*m*=−0.02, −0.01–−0.02, one-tailed *p*<0.001; Figure **2e**). These findings confirm the strong influence of spatial attention on long-term memory.

We predicted that long-term memory would also reflect trial-by-trial fluctuations of sustained attention, as observed in Experiment 1a. To control for whether items were cued to be spatially attended, we repeated the binning analysis within each condition (cued, uncued, and neutral; Figure **2c**). We calculated the slope that related long-term memory bin number to memory encoding, i.e., working memory response error (Figure **2d**). We observed a reliably positive relationship between long-term memory and working memory, for cued items (*m*=2.43, 1.74–3.29; one-tailed *p*<0.001), uncued items (*m*=4.04, 2.91–5.29; one-tailed *p*<0.001), as well as neutrally cued items (*m*=3.80, 2.94–4.90; one-tailed *p*<0.001). This replicated the finding that trial-by-trial fluctuations appeared to influence long-term memory encoding for items that were cued to be spatially attended and items that were not.

Finally, we examined whether these results replicated the finding that sustained attention fluctuations operate in an item-general manner. We examined long-term memory for items that were cued but unprobed and related it to long-term memory for items that were uncued but probed from the same display (Figure **2d**). Indeed, long-term memory for the unprobed items was correlated with long-term memory bins, as determined by separate items from the same display (*m*=1.81, 0.33–3.51, one-tailed *p*=0.01). We also replicated this finding with unprobed items in the neutrally cued blocks (*m*=2.24, 0.37–4.22, one-tailed *p*=0.01). This replicates the finding from Experiment 1a that fluctuations of sustained attention broadly impact memory for the entire display.

#### Recognition memory

We designed this experiment to measure long-term location memory via continuous report, in order to obtain a sensitive measure of memory fidelity. However, one possible concern is that participants sometimes could have been relying on their memory for the spatial cue, not the item itself. Therefore, in the long-term memory phase, we also measured item recognition memory. Indeed, overall recognition memory sensitivity was well above chance in Experiment 1 (*A′*=0.83; 0.80–0.85; one-tailed*p*<0.001 vs. chance=0.5).

We examined whether recognition memory correlated with performance at encoding. That is, we conducted a subsequent memory analysis for items that were later recognized versus not. Indeed, working memory response error was lower for items that were later recognized (*d*_recog_=17.39, 15.61–19.42; *d*_unrecog_=24.09, 21.29–27.22; one-tailed *p*<0.001). This shows a general correlation between memory encoding performance and later item recognition.

Item recognition was related to spatial and sustained attention. To examine the effect of spatial attention, we calculated the proportion of items that had been initially cued, separately for items that were later recognized vs. not. A greater proportion of items that were later recognized were initially cued (*q*_recog_=0.77, 0.76–0.78; *q*_unrecog_=0.72, 0.71–0.73; one-tailed *p*<0.001). To examine sustained attention, we calculated the working memory response error, separately for items that were later recognized vs. not. Indeed, working memory response error was lower for items that were later recognized, for cued items (*d*_recog_=11.83, 10.40–13.45; *d*_unrecog_=17.04, 14.63–19.83; one-tailed *p*<0.001) and uncued items (*d*_recog_=31.16, 27.40–35.81; *d*_unrecog_=35.90, 31.16–41.29; one-tailed *p*=0.006). Finally, we also observed that long-term item recognition memory reflected item-general fluctuations of sustained attention across displays: The memory hit rate for items that were unprobed was higher when the item from the same display was recognized (*h*_recog_=0.18, 0.15–0.20; *h*_unrecog_=0.13, 0.11–0.16; one-tailed *p*<0.001). That is, we observed a correlation in recognition memory performance between items from the same display.

In sum, recognition memory corroborated the findings from long-term location memory obtained via continuous report. These findings provide evidence for two distinct attentional signatures, sustained and spatial attention, which critically influence long-term memory.

### Discussion

This experiment demonstrates a robust influence of two distinct attentional factors, spatial attention and sustained attention, on the encoding of visual information into long-term memory. Long-term memory reflected whether participants were spatially attending the memoranda, as cued items were better remembered. Long-term memory also reflected putative fluctuations of sustained attention, exhibited as trial-by-trial fluctuations in memory encoding quality that affected memory for both spatially attended and spatially unattended items. We also observed additional evidence of sustained attention via trial-by-trial fluctuations of long-term memory for items from the same display. In sum, this experiment provided evidence of the composite model of attention and long-term memory.

In Experiment 2, we will extend these findings by measuring EEG activity while participants performed a similar task. This will provide the opportunity to identify neural signals that tracked fluctuations in sustained attention and the current locus of covert spatial attention. To anticipate the findings, differences in sustained attention are detectable based on neural activity even prior to stimulus onset, and this appears to be separate from fluctuations in the quality of spatial orienting of attention.

## Experiment 2

The goal of this experiment was to characterize the neural signals of sustained and spatial attention that predict long-term memory. We collected eye-tracking and EEG data while participants performed the task from Experiment 1a.

### Methods

#### Participants

Forty-two adults (23 female; mean 23.5 years) completed Experiment 2 for $60 payment ($15/hour). A larger number of participants was chosen so as to have an adequate sample size after excluding participants who had excessive EEG or eye artifacts (six participants who all had fewer than half of the trials remaining after artifact rejection of the prestimulus period) or problems with EEG or eye-tracking equipment during the recording session (six participants). The final sample size was thirty participants. These exclusion criteria were determined a priori and are consistent with prior studies from our laboratory.

#### Apparatus

Participants were seated approximately 75 cm from a LCD monitor (120-Hz refresh rate) in a shielded booth.

#### Stimuli & procedure

Same as Experiment 1a.

#### Eye tracking

We monitored gaze position using a desk-mounted infrared eye tracking system (EyeLink 1000 Plus, SR Research, Ontario, Canada). Gaze position was sampled at 1000Hz, and head position was stabilized with a chin rest.

#### EEG recording

We recorded EEG activity using 30 active Ag/AgCl electrodes mounted in an elastic cap (Brain Products actiCHamp, Munich, Germany). We recorded from International 10/20 sites: FP1, FP2, F3, F4, F7, F8, Fz, FC1, FC2, FC5, FC6, C3, C4, Cz, CP1, CP2, CP5, CP6, P3, P4, P7, P8, Pz, PO3, PO4, PO7, PO8, O1, O2, and Oz. Two electrodes were placed on the left and right mastoids, and a ground electrode was placed at position FPz. All sites were recorded with a right-mastoid reference, and were re-referenced offline to the algebraic average of the left and right mastoids. Eye movements and blinks were recorded with passive electrodes using horizontal and vertical EOG. Data were filtered online (0.01 to 250 Hz) and were digitized at 1000 Hz using BrainVision Recorder.

#### Artifact rejection

We extracted data relative to the onset of spatial attention cues (−300 to 1500 ms relative to cue onset). Using an automatic algorithm, trials were examined for EEG artifacts (amplifier saturation, excessive muscle noise, and skin potentials), and EOG & eye-tracking ocular artifacts (blinks and saccades exceeding 0.5° from fixation). We also manually inspected all trials. Participants were excluded if fewer than half of the trials remained after discarding those with artifacts. On average, 295 (77%) trials remained per participant after artifact rejection.

#### Multivariate classification

For all artifact-free trials, we used multivariate pattern classifiers to predict long-term memory. We decomposed EEG event-related potentials into multifrequency oscillatory bands (4–7 Hz, 8–12 Hz, 13–16 Hz, 16–20 Hz, 20–25 Hz, 25–30 Hz), to encompass the wide variety of frequency bands have been implicated in prior work investigating long-term memory. We first band pass filtered the data and applied the Hilbert transform. To equate for power differences across frequency bands without removing sustained pre-stimulus signals, we demeaned the signal based on the global average power within each band. We labeled trials according to whether they were accurately or inaccurately remembered, based on the median absolute response error in the long-term memory phase. To eliminate spatial imbalances, we calculated the median per quadrant. Therefore, accurate and inaccurate trials contained the same number of items per quadrant.

We trained a multivariate classifier using L2-penalized logistic regression (C=1) with the scikit-learn package in python. We split trials into two equal sized bins (training and test). The input data was the power at each frequency at each electrode (100 ms time window). We averaged trials within bins and then repeated this entire procedure for 10 ms timesteps. We further repeated the assignment of trials to bins 1,000 times for each participant. We conducted statistical analyses across participants to compare classification accuracy to theoretical chance (50%) as well as a shuffled null, for which we permuted the labels.

To decode spatial attention, we repeated the general procedure of multivariate pattern analysis. We decomposed EEG event-related potentials into alpha (8–12 Hz) power. We chose to focus our analysis on the alpha frequency band a priori based on extensive work showing classification of spatial attention in this band (Foster et al., 2017). We labeled all accurate and inaccurate trials according to the cued quadrant. Then we split accurate and inaccurate trials each into two bins. We downsampled and then averaged trials within each bin×quadrant. We trained a classifier based on a mixed training set, using one bin of accurate and one bin of inaccurate trials. We tested the classifier separately on the held out bin of accurate or inaccurate trials. We further repeated the assignment of trials to bins 1,000 times for each participant. We conducted statistical analyses across participants to compare classification accuracy to theoretical chance (25%) as well as a shuffled null, for which we permuted the labels. Code that describes the entire multivariate decoding procedure is available online.

#### Statistics

Same as Experiment 1.

### Results

#### Behavioral results

We again operationalized long-term memory as absolute response error (*d_ltm_*=61.10°, 95% CIs 56.14–65.92; Figure **3a**). We replicated our findings from Experiment 1, that general correlation between memory encoding and long-term memory. We observed a reliably positive slope relating long-term memory (i.e., bin number) to memory encoding via working memory response error (*m*=2.21, 1.70–2.89; one-tailed *p*<0.001; Figure **3b**). We further disentangled how long-term memory reflected both spatial and sustained attention:

**Figure 3.**
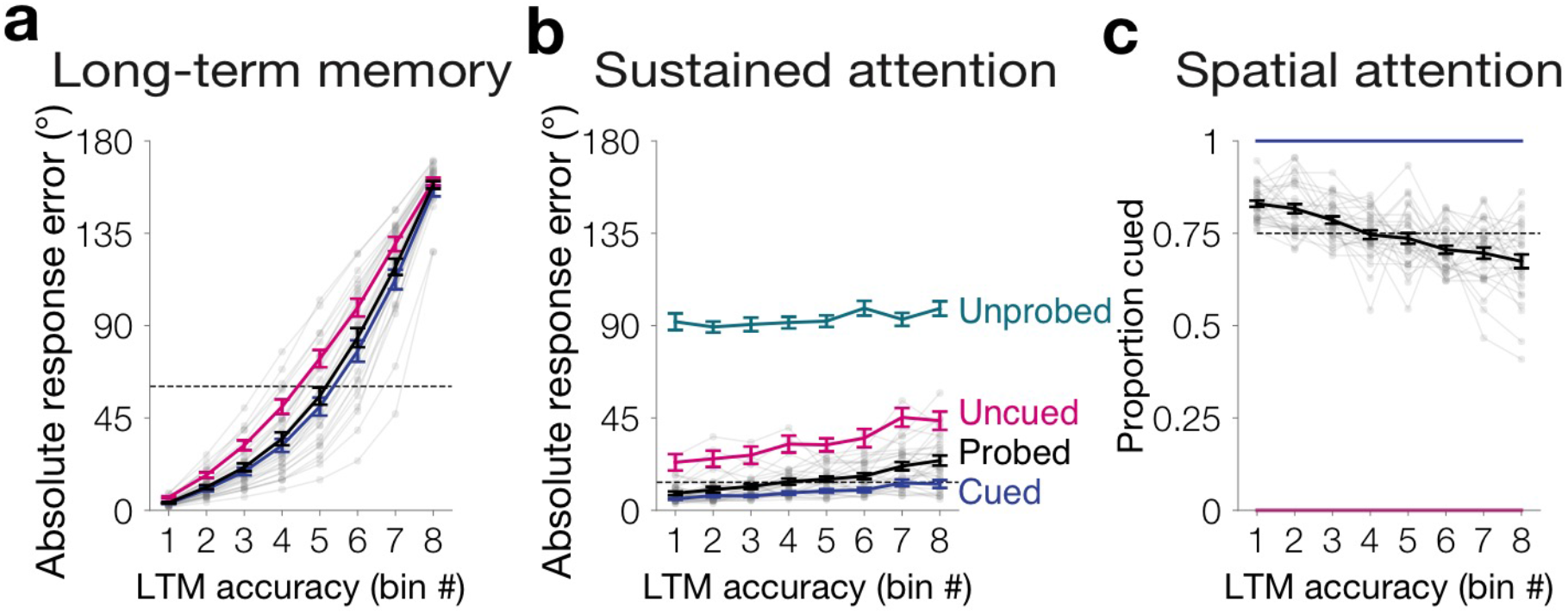
Experiment 2 behavioral results. **a** Long-term location memory variability binned across trials. Absolute response error (0–180°) was sorted and binned into octiles within participant for all probed items (black line). To equate for whether an item was cued to be attended, we also repeated all analyses separately for cued items (blue), and uncued items (pink). The median long-term memory response error for all probed items is depicted as a dashed black line. **b** Memory outcomes reflect trial-by-trial fluctuations of sustained attention. Memory encoding success was operationalized as absolute response error (0–180°) for items within each bin. For uncued (pink), probed (black), and cued (blue) items, absolute response error was obtained during the working memory phase. The median working memory response error for probed items is depicted as a dashed black line. Unprobed items (teal) were not tested in the working memory phase. Therefore, absolute response error for these items was obtained during the long-term memory phase. The slope of each line is positive across the bins (*p*s<0.05). **c** Memory outcomes reflect trial-by-trial differences of spatial attention. In each bin, we calculated the proportion of trials that had been cued. Within all probed items (black), the proportion of trials that were cued decreases across the eight bins (*p*<0.001). The dashed line represents the mean proportion cued (0.75). Cued items (blue) and uncued items (pink) controlled for whether an item was cued to be attended (100% and 0% cued, respectively). Error bars depict the standard error of the mean. Data from each participant from all probed items with a single cue are overlaid in small gray dots connected with lines.

We replicated the finding that long-term memory reflected spatial attention. Items that were cued to be spatially attended were better remembered (*d_cued_*=57.69°, 52.35–62.98; *d_uncued_*=71.34°, 66.19–75.86; one-tailed *p*<0.001). This relationship between spatial attention and long-term memory was further quantified as a negative slope of the proportion cued across bins (*m*=−0.023, −0.031–−0.017; one-tailed *p*<0.001; Figure **3c**).

We also replicated the finding that long-term memory appeared to reflect trial-by-trial fluctuations in sustained attention within each condition (Figure **3b**). We observed a positive slope relating long-term memory to encoding for cued items (*m*=1.09, 0.74–1.77, p<0.001) and uncued items (*m*=3.18, 2.27–4.26; one-tailed *p*<0.001). Furthermore, we examined the long-term memory response error per bin for the items that had been cued but were not probed from the uncued trials. We observed a positive slope relating long-term memory for the uncued items (those that had been probed in the working memory phase) to long-term memory for the unprobed items from the same display (*m*=1.05, −0.23–2.17; one-tailed *p*=0.045). In sum, behavioral evidence replicated the observation that long-term memory reflects trial-by-trial fluctuations of spatial and sustained attention at encoding.

#### EEG

We observed from behavioral evidence the importance of trial-to-trial fluctuations. However, these behavioral findings were unable to specify when and how these fluctuations emerged. By examining EEG activity that tracks the subjects’ current attentional state, we developed a classifier to investigate pre-stimulus fluctuations of sustained attention. We tested whether ongoing neural activity could predict long-term memory success even prior to item presentation. Given that participants had to sustain attention over an extended 2–4 second postcue interval, we targeted this post-cue and pre-stimulus time window for analyses. We used multivariate classification to decode whether a trial was remembered *accurately* or *inaccurately*, relative to the median long-term memory response error per quadrant (Figure **4a**). Indeed, we reliably decoded long-term memory accuracy based on EEG patterns in the time window following the cue (mean accuracy=57.23%, 95% CIs 52.87–62.58%; *n*=30, one-tailed *p*<0.001, chance=50%; *t*=500–100 ms; Figure **4b**). That is, we could predict whether an upcoming item would be better remembered, even before it appeared. These decoding results are consistent with trial-by-trial fluctuations of sustained attention that occur prior to stimulus presentation and influence memory encoding.

**Figure 4.**
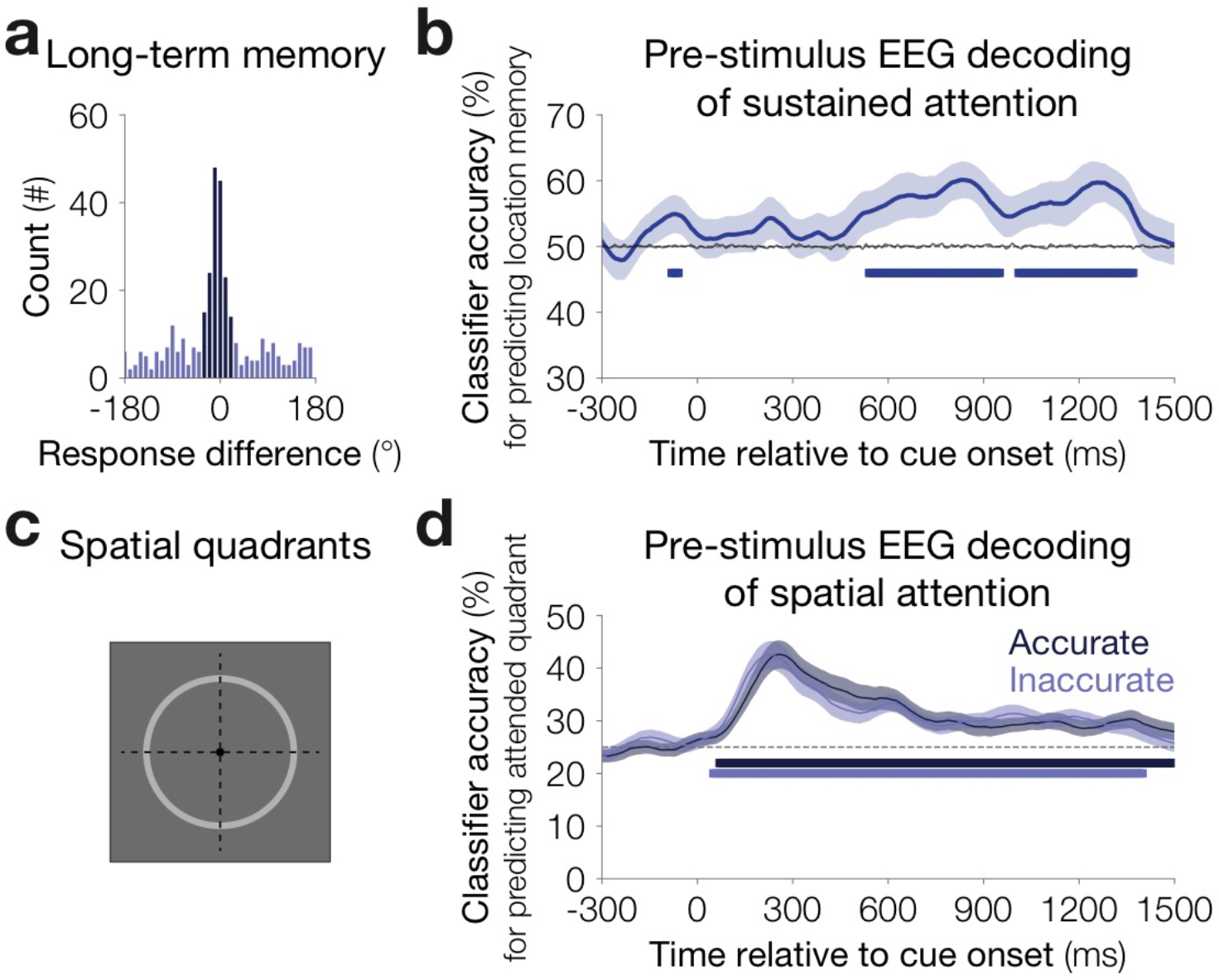
Multivariate decoding of attentional processes from EEG data. **a** Long-term memory accuracy for a representative participant. A histogram of response differences across trials, the angular distance from the original image location (−180 to 180°). Labels for the multivariate pattern classifier reflected the median absolute response error per quadrant, either *accurate* (dark blue) or *inaccurate* (light blue). **b** Pre-stimulus EEG decoding of sustained attention. Long-term memory accuracy can be predicted prior to stimulus onset. A multivariate classifier was trained to predict long-term location memory *(accurate* vs. *inaccurate)* based on multi-frequency EEG patterns and tested on held out trials. Average classification accuracy is depicted as the solid blue line, the gray line depicts empirical chance after shuffling the labels 1,000 times. The shaded area is the standard error of the mean. Blue squares highlight timepoints for which classification accuracy is above chance (*p*<0.05). **c** Spatially attended quadrants. The display visually depicts the labels provided to the labels spatial attention classifier. Trials were split according the cued quadrant (1–4), four quadrants illustrated by dashed lines. **d** Pre-stimulus EEG decoding of longterm memory is not explained by differences in spatial attention. The cued quadrant was reliably decoded from both accurately and inaccurately remembered items. A multivariate classifier was trained to predict cued location and tested on held out *accurate* or *inaccurate* trials. Average classification accuracy is depicted as solid lines for accurate trials (dark blue) and inaccurate trials (light blue). Squares depict timepoints in which classification accuracy for either condition is above chance. The shaded area is the standard error of the mean.

The behavioral evidence suggested that sustained attention is distinct from spatial attention, as it broadly impacts both cued and uncued items from the same display. Therefore, we predicted that fluctuations of sustained attention should be independent from fluctuations of spatial attention. If this is the case, then our classifier of sustained attention should be robust, even when we collapse data across all spatial locations. Alternatively, fluctuations of sustained attention could have reflected general task disengagement. If so, fluctuations of sustained attention would predict worse spatial attention during low moments of sustained attention. To explore these possibilities, we investigated whether we could decode the cued quadrant (Figure **4c**). During inaccurately remembered trials, we reliably decoded the cued position (mean accuracy=31.82%, 95% CIs 29.99–33.98%; *n*=30; one-tailed *p*<0.001, chance=25%; *t*=0–1500 ms; Figure **4d**). We also reliably decoded the cued position during accurately remembered trials (mean accuracy=31.99%, 95% CIs 30.17–34.90%; *n*=30; one-tailed *p*<0.001, chance=25%; *t*=0– 1500 ms; Figure **4d**). Critically, there was no reliable difference in spatial attention decoding between accurate and inaccurate trials (two-tailed *p*=0.76). These decoding results suggest that the ability to predict long-term memory during pre-stimulus windows is not driven by differences in spatial attention.

#### Recognition memory

Finally, we examined item recognition memory behavior. Overall recognition memory was above chance for all probed items (*A*′=0.87; 0.85–0.89; one-tailed *p*<0.001 vs. chance=0.5). Recognized items exhibited a lower absolute response error in the working memory phase (*d*_recog_=14.30, 12.31–17.35; *d*_unrecog_=16.48, 14.16–19.28; one-tailed *p*<0.001), suggesting that recognition memory reflected trial-by-trial fluctuations in memory encoding. However, this could have been in part driven by the effect of spatially attending an item (*q*_recog_=0.76, 0.75–0.78; *q*unrecog=0.73, 0.71–0.74; one-tailed *p*<0.001). Therefore, we controlled for whether items were spatially attended, and replicated the effects of trial-by-trial fluctuations for cued items (*d*_recog_=8.73, 7.59–10.43; *d*_unrecog_=10.13, 8.4–12.76; one-tailed *p*=0.01). We did not replicate the effect of sustained attention on recognition memory for uncued items (*d*recog=32.01, 25.95–40.75; *d*_unrecog_=33.74, 27.70–41.24; one-tailed *p*=0.20) or unprobed items (*h*_recog_=0.16, 0.13–0.19; *h*_unrecog_=0.15, 0.12–0.18; one-tailed *p*=0.15). Though item recognition results are largely consistent with results obtained from continuous report, these findings also suggest that continuous report may provide a more sensitive assay of how spatial and sustained attentional factors influence long-term memory.

### Discussion

Experiment 2 replicated and extended the behavioral findings from Experiment 1, with concurrent eye-tracking to ensure spatial attention was maintained covertly. We demonstrated that pre-stimulus multivariate EEG patterns covaried with long-term memory. These results revealed a multivariate EEG representation of sustained attentional state that preceded stimulus onset and related to long-term memory.

Furthermore, these behavioral and neural findings supported the composite model of attention and long-term memory, as fluctuations of sustained attention were distinct from spatial attention. These results also rule out the interpretation that trial-by-trial fluctuations of sustained attention reflect moments during which participants completely disengaged from the task: EEG activity showed that subjects maintained covert spatial attention at the cued position, even when the cued stimulus was later inaccurately remembered. Thus, multivariate analyses of EEG data suggest that fluctuations in sustained attention can be distinguished from the waxing and waning of spatial attention at cued positions, and do not reflect episodes of global disengagement with the task.

## Experiment 3

Thus far, we have focused primarily on recall of the spatial position of the memoranda. Experiment 3 examined whether distinct attentional subcomponents also influenced long-term memory for non-spatial features. We adjusted the paradigm to test memory for a continuous but non-spatial feature of the item, namely, color. We hypothesized that trial-to-trial fluctuations in sustained and spatial attention would distinctly predict long-term memory for color.

### Methods

#### Participants

In Experiments 3a and 3b, a combined total of fifty-two adults participated for University of Chicago course credit or $20 payment ($10/hour). In Experiment 3a, twenty-five adults (15 female, mean age = 23.0 years) participated. One participant was excluded due to errors during data collection, resulting in a final sample size of 24. In Experiment 3b, twentyseven adults (20 female, mean age = 20.1 years) participated. One participant was excluded due to errors during data collection, resulting in a final sample size of 26.

#### Apparatus

Same as Experiment 1b.

#### Stimuli

A subset of the real-world object images from Experiments 1–2 selected based on the relative uniqueness of the shape outlines were manipulated to be a one-dimensional color mask. The color of each image was sampled randomly from a 360° HSV space, with saturation and value of 1, and remapped to RGB values for presentation in Psychopy.

#### Procedure

In the memory probes, participants reported the color memory for each item instead of its spatial location (Figure **5a**). The item, colored dark gray, appeared at the center of the screen, surrounded by a color wheel. The color wheel was randomly oriented for each trial, and this random orientation was held consistent between working memory and long-term memory tests. Based on piloting, we reduced the working memory array to 2 items, which were separated by a minimum distance of 40° in color space. In the long-term memory phase, participants made source memory judgments only for the probed images from the working memory phase, both cued or uncued. In Experiment 3a, participants completed 24 blocks of 16 trials, and participants completed on average 368 of the maximum 384 trials (95.83%), ranging from 272 (70.83%) to 384 (100%). To boost color long-term memory performance in Experiment 3b, we eliminated the recognition memory judgments and reduced the block length. In Experiment 3b, participants completed 48 blocks of 8 trials and all participants completed all 384 trials.

**Figure 5.**
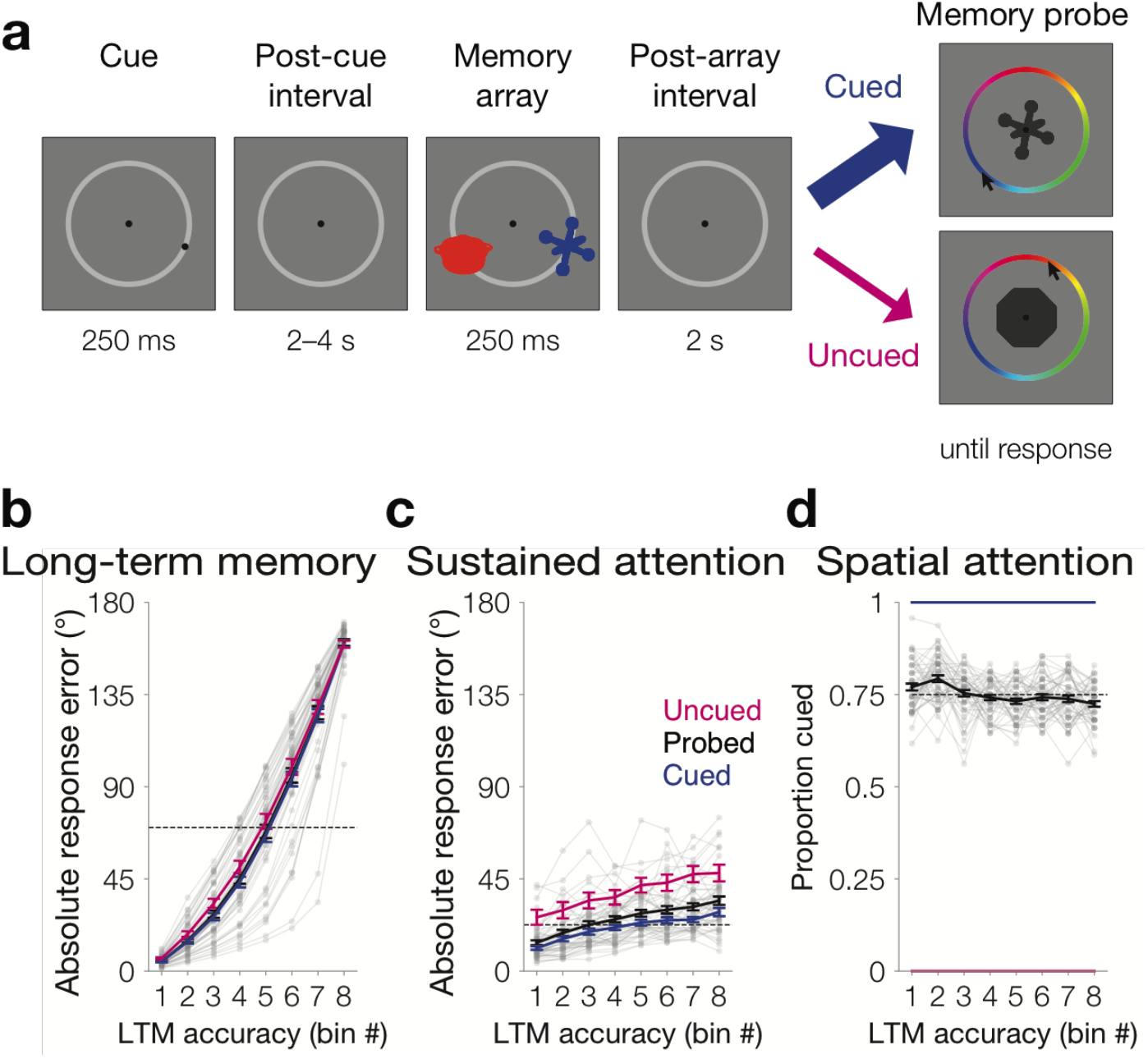
Experiment 3a&b color memory results. **a** Task design of the working memory phase. Each trial was comprised of a brief peripheral spatial cue, a blank pre-stimulus delay, a memory array of two trial-unique object pictures, and a blank retention interval. The object pictures were filled with a color from a continuous color wheel. For the memory probe, one of the items (cued: 75%; uncued 25%) reappeared at the center in dark gray, and participants reported its original color by clicking along the wheel. **b** Long-term memory variability binned across trials. Color memory absolute response error (0–180°) was sorted and binned into octiles within participant for all probed items (black line). To equate for whether an item was cued to be attended, we also repeated all analyses separately for cued items (blue) and uncued items (pink). The median longterm memory response error for all probed items is depicted as a dashed black line. **c** Memory outcomes reflect trial-by-trial fluctuations of sustained attention. Memory encoding success was operationalized as absolute response error for items within each bin (0–180°). For uncued (pink), probed (black), and cued (blue) items, absolute response error was obtained during the working memory phase. The median working memory response error for probed items is depicted as a dashed black line. **d** Memory outcomes reflect trial-by-trial differences of spatial attention. In each bin, we calculated the proportion of trials that had been cued. For all probed items (black), the proportion of trials that were cued decreases across the eight bins (*p*<0.001). The dashed line represents the mean proportion cued (0.75). Cued items (blue) and uncued items (pink) controlled for whether an item was cued to be attended (100% and 0% cued, respectively). Error bars depict the standard error of the mean. Data from each participant from all probed items with a single cue are overlaid in small gray dots connected with lines.

#### Behavioral analysis & Statistics

Same as Experiments 1–3.

### Results

#### Experiment 3a

The goal of this experiment was to replicate the findings from Experiments 1 and 2 and extend them to the color dimension. We operationalized long-term memory as absolute response error from the original color (*d_ltm_*=71.57°, 95% CIs 63.97–76.59). For each participant, we sorted and binned trials according to their long-term color memory performance (Figure **5b**). We observed that long-term color memory bins reflected trial-by-trial fluctuations in color working memory encoding, as a positive slope that related working memory to long-term memory across bins (*m*=2.35, 1.93–2.75; one-tailed *p*<0.001; Figure **5c**).

We next examined whether long-term color memory also reflected distinct factors of spatial and sustained attention. To examine the influence of sustained attention, we observed that long-term color memory was enhanced by spatial attention (*d_cued_*=70.87°, 63.24–75.88; *d_uncued_*=73.68°, 65.85–79.14; one-tailed *p*=0.006). Furthermore, the proportion of items that had been cued declined across bins (*m*=−0.005, −0.008–−0.002; one-tailed *p*=0.001; Figure **5d**).

To examine the influence of sustained attention, we repeated the binning analysis within each cueing condition (Figure **5c**). We observed that long-term memory bin was positively related to working memory response error for cued items (*m*=1.93, 1.54–2.33, p<0.001) and uncued items (*m*=2.86, 1.82–3.82; one-tailed *p*<0.001). In sum, behavioral evidence confirms that trial-by-trial fluctuations are critical in understanding long-term color memory.

#### Experiment 3b

The goal of this experiment was to replicate the findings of Experiment 3a, while boosting the long-term memory phase color memory performance by reducing the number of trials per block (*d_ltm_*=64.51°, 95% CIs 58.95–68.91). We replicated the binning analysis based on long-term color memory performance (Figure **5b&c**). We again observed that long-term color memory could be explained by trial-to-trial fluctuations in color working memory performance (*m*=3.16, 2.52–3.97; one-tailed *p*<0.001). We investigated whether this general correlation between working memory and long-term memory reflected key signatures of spatial and sustained attention.

We replicated the influence of spatial attention on long-term color memory. We observed that long-term color memory was more accurate for cued items (*d_cued_*=62.94°, 57.72–67.28; *d_uncued_*=69.21°, 62.35–74.79; one-tailed *p*<0.001). In addition, the proportion of items that had been cued declined across bins (*m*=−0.010, −0.016–−0.005; one-tailed *p*<0.001; Figure **5d**).

To investigate the influence of trial-by-trial fluctuations of sustained attention on longterm memory for color, we again repeated the binning analysis within each cueing condition (Figure **5b**). We still observed a reliably positive slope between long-term memory bin and working memory response error for cued items (*m*=2.55, 2.00–3.27, p<0.001) and uncued items (*m*=3.57, 2.60–4.76; one-tailed *p*<0.001). This, once again, demonstrates that trial-by-trial fluctuations are critical for long-term color memory.

#### Recognition memory

We only collected long-term recognition memory ratings in Experiment 3a. In this experiment, overall recognition memory sensitivity was well above chance for all probed items (*A′*=0.78, 0.74–0.82; one-tailed *p*<0.001 vs. chance=0.5). We observed a general relationship between item recognition and working memory performance, as working memory response error was lower for recognized vs. unrecognized items (*d*_recog_=23.52, 19.95–29.51; *d*_unrecog_=29.51, 25.67–34.64; one-tailed *p*<0.001).

We conducted a subsequent memory analysis to examine whether spatial and sustained attention related to item recognition. We did not observe a reliable effect of spatial attention, as recognized trials were not more likely to have been cued (*q*_recog_=0.76, 0.75–0.78; *q*_unrecog_=0.75, 0.74–0.75; one-tailed *p*=0.08). However, we did observe that trial-by-trial fluctuations of sustained attention influenced item recognition memory for cued items. Working memory absolute response error was lower for items that were later recognized vs. not, for cued items (*d*_recog_=19.50, 16.66–24.47; *d*_unrecog_=23.69, 20.29–28.18; one-tailed *p*<0.001) and uncued items (*d*_recog_=37.76, 29.59–51.29; *d*_unrecog_=46.14, 38.02–58.34; one-tailed *p*<0.001).

### Discussion

These findings verified that attention also influences memory for a non-spatial feature. Indeed, both sustained and spatial attention distinctly related to long-term color memory. These results extend the finding that sustained attention fluctuations are a broad and general influence for memories that can be clearly dissociated from the effects of spatial attention.

## Individual differences

The goal of this study was primarily to explore attention and long-term memory within individuals. However, we also examined between-subject variations in long-term memory performance, and the underlying attentional factors that drove those differences. We collapsed across data from all experiments to boost the number of subjects (*n*=127). We were interested in the exploring the degree to which individual differences in long-term memory reflected individual differences in sustained or spatial attention.

### Methods

#### Participants

We collapsed across data from all participants in Experiments 1–3 (*n*=127) to maximize the number of subjects.

#### Behavioral analysis

We examined measures of sustained and spatial attention from the binning analysis of Experiments 1–3. For each participant, we computed a measure of sustained attention as the slope that related long-term memory bin number to working memory absolute response error. We computed the slope separately for cued and uncued items and averaged across conditions for a single measure of sustained attention. We also computed a measure of sustained attention as the slope that related long-term memory bin number to the proportion uncued (1-proportion cued per bin). We examined how measures of sustained and spatial attention related to average long-term memory absolute response error. We also examined how measures of sustained and spatial attention related to each other.

To examine the reliability of these measures of sustained attention, we conducted a downsampling analysis: We randomly sampled 50% of trials to calculate a downsampled measure of sustained attention and repeated this downsampling procedure to calculate a downsampled measure of spatial attention. To ensure that experiment-specific effects were not driving the individual differences, we also z-scored sustained attention, spatial attention, and long-term memory within experiment and repeated the analyses.

#### Statistics

Descriptive statistics are reported as the mean and 95% confidence interval (CI) of the bootstrapped distribution. Correlations and partial correlations were computed using the non-parametric Spearman’s rank-order correlation function. Linear models were fit in R, and we report the adjusted *r*^2^ values. F statistics evaluated different model fits. For the downsampling analyses, we report the 95% confidence intervals across downsampling iterations.

### Results

We calculated a measure of *sustained attention* for each participant in our study, as the slope of the working memory response error across bins. A more positive slope reflects a stronger influence of sustained attention on memory. That is, items that were encountered in a more advantageous sustained attentional state were better remembered (relative to items in a disadvantageous attentional state). To eliminate any influence of spatial cues, we calculated the slope separately within condition (cued and uncued) and averaged across conditions. If sustained attention benefitted overall memory performance, then individuals who demonstrated a stronger influence of sustained attention (i.e., more positive slope) would also demonstrate better longterm memory performance. Indeed, there was a reliable relationship between sustained attention and long-term memory response error (β_sust_=−2.68, −3.90–−1.46; *r*^2^=0.12; *p*<0.001; Figure **6a**). That is, individual differences in sustained attention correlated with long-term memory performance.

**Figure 6.**
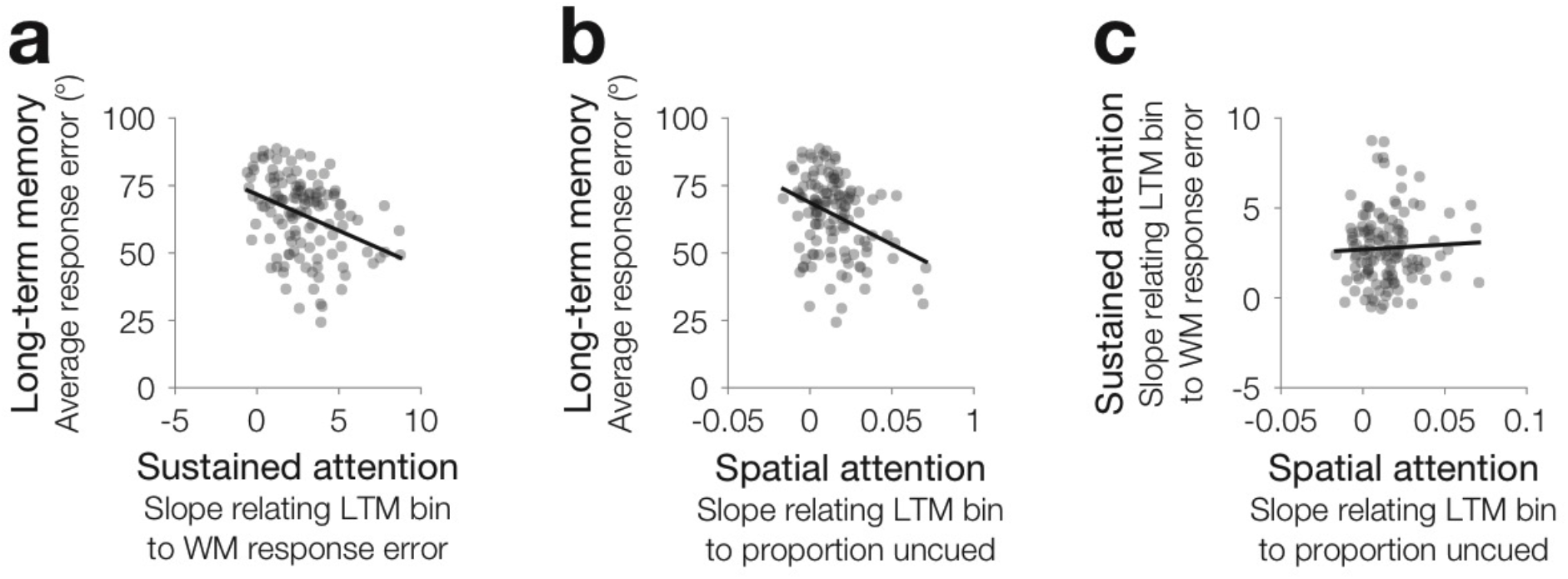
Individual differences. **a** Individual differences in sustained attention relate to long-term memory. For each participant, we calculated a measure of sustained attention, as the slope that related long-term memory (LTM) bin to working memory (WM) response error. A more positive value of sustained attention reflects a stronger influence of sustained attention on long-term memory. We observed that sustained attention was negatively correlated with absolute response error in the long-term memory phase (*p*<0.001). That is, a stronger influence of sustained attention correlates with better overall long-term memory. **b** Individual differences in spatial attention relate to long-term memory. For each participant, we calculated a measure of spatial attention, as the slope that related long-term memory (LTM) bin to the proportion of uncued items per bin. A more positive value of spatial attention reflects that long-term memory was especially related to whether an item was spatially cued. We observed a stronger influence of spatial attention was absolute response error in the long-term memory phase (*p*<0.001). That is, a stronger influence of spatial attention correlates with better overall long-term memory. **c** Individual differences in sustained and spatial attention are unrelated. We examined whether these measures of sustained attention was correlated with spatial attention across individuals. There was no reliable relationship between sustained and spatial attention. Data from all participants in Experiments 1, 2, and 3, are overlaid in gray dots. The linear relationship is depicted with a black line.

We also calculated a measure of *spatial attention* for each participant in our study, as the slope of the proportion uncued across bins. A more positive slope reflects a stronger influence of spatial attention. That is, items that were cued were better remembered (relative to items that were uncued). If spatial attention benefitted overall memory performance, then individuals who demonstrated a stronger influence of spatial attention (i.e., more positive slope) would also demonstrate better long-term memory performance. Indeed, there was reliable relationship between spatial attention and long-term memory response error (β_spatial_=−310, −459–−161, *r*^2^=0.11, *p*<0.001; Figure **6b**). That is, individual differences in spatial attention correlated with long-term memory performance.

The critical question is whether these two signals, sustained and spatial attention, reflect separate influences on long-term memory. If so, we would predict that they are uncorrelated with each other and explain a unique portion of the variance. We found that individual differences in spatial attention and sustained attention were uncorrelated with each other (*r*=0.02, *p*=0.81; Figure **6c**). In addition, they explained unique variance in individual differences in long-term memory: The partial correlation between sustained attention and long-term memory (*r*=−0.38, *p*<0.001) and the partial correlation between spatial attention and long-term memory (*r*=−0.29, *p*<0.001) were both reliable. The model using both signals as predictors explained more variance in individual differences in long-term memory (adjusted *r*^2^=0.23) than a model that just included sustained attention (F_(1,125)_=17.63, *p*<0.001) or spatial attention (F_(1,125)_=19.70, *p*<0.001). In sum, individual differences for sustained attention and spatial attention distinctly influence long-term memory.

We further inspected these results to examine the reliability of the measures of sustained and spatial attention. To obtain a measure of sustained attention, we collapsed across both the cued and uncued conditions. Across individuals, the measure of sustained attention obtained from cued and uncued conditions were correlated with each other (*r*=0.43, *p*<0.001). Furthermore, we ran a downsampling analysis where we randomly selected 50% of trials per participant and reran the binning analyses using these trials. Both sustained attention and spatial attention remained reasonably stable across downsampling iterations (*m*_sust_=2.72, 2.54–2.89; *m*_space_=0.015, 0.013–0.016). In addition, the relationship between sustained attention and spatial attention with long-term memory remained negative across downsampling iterations (β_sust_=−1.91, −2.43–−1.40; β_spatial_=−219, −297–−153). Thus, these measures of sustained and spatial attention appear reliable.

Finally, to ensure that these correlations did not arise from spurious differences in the experiments, we repeated all analyses after z-scoring across participants within each experiment. Even after z-scoring, sustained attention reliably predicted memory (β_sust_=−5.61, −7.97–−3.25; *r*^2^=0.14; *p*<0.001), and spatial attention reliably predicted memory (β_spatial_=−3.37, −5.85–−0.88, *r*^2^=0.05, *p*=0.008). Sustained and spatial attention were still uncorrelated (*r*=0.06, *p*=0.50). In addition, the model that included both sustained and spatial attention better predicted long-term memory (adjusted *r*^2^=0.18) than a model that just included sustained attention (F_(1,125)_= 6.16, *p*=0.014) or a model that just included spatial attention (F_(1,125)_=20.95, *p*<0.001). Thus, effects across experiments did not appear to drive the individual differences.

### Discussion

We examined the influence of sustained and spatial attention on long-term memory performance across participants in the three experiments. These findings revealed that sustained and spatial attention predict distinct variance in long-term memory performance.

## General Discussion

Over three experiments, we found that spatial and sustained attention had robust and distinct effects on long-term memory. Behaviorally, long-term memory was predicted from whether spatial attention was correctly oriented and sustained attentional state was advantageous. Spatial attention was manipulated by pre-cues, and fluctuations in sustained attention were observed via trial-to-trial variations in working memory performance. Neurally, spatial and sustained attentional states were decoded from multivariate analyses of EEG data prior to stimulus presentation. EEG measures of sustained attention were distinct from spatial attention, as they generalized across spatial positions. Furthermore, sustained attention differences did not reflect general task disengagement, as the cued location was equally well decoded during lapses of sustained attention. We provide evidence that sustained and spatial attention can have orthogonal and even parallel impacts on memory. In sum, memory failures can be attributed to failures of distinct spatial or sustained attentional processes, as shown by these behavioral results, neural analyses, and individual differences.

This work provides critical support for a composite model of attention and long-term memory. That is, distinct attentional subcomponents can each explain what we fail to remember. We can conceptually illustrate the consequences of distinct attentional processes with an addendum to the traditional spotlight analogy: If spatial attention reflects the location of the spotlight on the stage, sustained attention corresponds to the number of audience members who are awake. Critical to this analogy is the idea that sustained and spatial attention can each vary independently. That is, even when attention is *oriented* correctly, it could be that sustained attention was *deployed* unsuccessfully. Furthermore, either the spotlight or the audience’s wakefulness can contribute to later memory of the performance, as suggested by the composite model of attention and long-term memory. Conversely, the unified model would suggest that sustained attention and spatial attention have largely overlapping influences (e.g., spotlight location and brightness), which we did not observe.

### Sustained vs. spatial attention

This study sheds light on how two distinct attentional processes impact encoding into long-term memory. Indeed, these findings converge with prior attention research that has shown spatial and sustained attention reflect distinct cognitive processes, both within and between individuals (Dowd & Golomb, 2019; J. Fan et al., 2005; Poole & Kane, 2009; Robison & Brewer, 2019). Our study makes the key contribution of extending the dissociable nature of sustained and spatial attention to the domain of long-term memory. Comprehensively characterizing the neural signatures of sustained attention and their key contribution to long-term memory will be important going forward. Traditionally, the relationship between attention and long-term memory has been investigated by manipulating topdown attention (Aly & Turk-Browne, 2017). Here, we argue that it is important to distinguish between multiple forms of attention, and specifically consider fluctuations of sustained attention, to understand how attention shapes memory.

### Sustained attention and long-term memory

This study also motivates further study of sustained attention, a growing area within recent attention research (deBettencourt et al., 2015, 2018, 2019; Esterman et al., 2013, 2014; Rosenberg et al., 2016). However, there remain many open questions about how sustained attention relates to long-term memory. Memory studies have often ignored the influence of attention, and rarely consider how sustained attentional state may dramatically impact memory formation (Adam & deBettencourt, 2019). One possibility is that previous demonstrations of pre-stimulus signals that predict memory are in fact tapping into sustained attentional state (Adam et al., 2015; Gruber & Otten, 2010; Guderian et al., 2009; Otten et al., 2006; Weidemann & Kahana, 2020). Future studies can develop a more complete understanding of the neural processes of sustained attention and how they are recruited for longterm memory.

### Arousal and memory

The distinct effects of sustained and spatial attention on long term memory encoding do not reflect general task engagement or global arousal. Even while sustained attention was lapsing, we decoded the spatially attended location equally well. That is, lower-level spatial representations are intact even while sustained attention lapses. Indeed, this finding is in line with previous study that observed lapses of sustained attention did not impact the precision of working memory representations (deBettencourt et al., 2019). Future work could more carefully characterize when and how general arousal contributes to measurements of sustained attention and predictions of long-term memory.

### Memory improvement

Finally, this work provides important suggestions for strategies to improve or worsen memory. Cueing attention enhances long-term memories at the attended location at the expense of memories elsewhere. In addition, harnessing sustained attention fluctuations may be another key for altering memory performance. Our findings suggest that advantageous moments of sustained attention broadly enhance memories for items distributed across spatial locations. Thus, targeting optimally attentive states or optimally attentive moments, or even improving general sustained attention abilities (deBettencourt et al., 2015), may be a key strategy for improving memory.

## Author contributions

All authors conceived of the study and contributed to the study design. M.T. deBettencourt and S.D. Williams collected and analyzed the data. M.T. deBettencourt wrote an initial draft of the manuscript, which all authors read and edited.

## Acknowledgements

This research was supported by National Institute of Health grant R01MH087214, the Office of Naval Research grant N00014-12-1-0972, and F32MH115597 (M.T.dB.). We thank A. Gale for assistance with data collection for Experiment 2, M. Bolouri for assistance with data collection for Experiment 3, and K.C.S. Adam, N. Hakim, and A. Tompary for comments on earlier versions of the manuscript.

